# Genetic evidence for natural selection in humans in the contemporary United States

**DOI:** 10.1101/037929

**Authors:** Jonathan Beauchamp

## Abstract

Recent findings from molecular genetics now make it possible to test directly for natural selection by analyzing whether genetic variants associated with various phenotypes have been under selection. I leverage these findings to construct polygenic scores that use individuals’ genotypes to predict their body mass index, educational attainment (EA), glucose concentration, height, schizophrenia, total cholesterol, and (in females) age at menarche. I then examine associations between these scores and fitness to test whether natural selection has been occurring. My study sample includes individuals of European ancestry born between 1931 and 1953 in the Health and Retirement Study, a representative study of the US population. My results imply that natural selection has been slowly favoring lower EA in both females and males, and are suggestive that natural selection may have favored a higher age at menarche in females. For EA, my estimates imply a rate of selection of about -1.5 months of education per generation (which pales in comparison with the increases in EA observed in contemporary times). Though they cannot be projected over more than one generation, my results provide additional evidence that humans are still evolving—albeit slowly, especially when compared to the rapid secular changes that have occurred over the past few generations due to cultural and environmental factors.

---

Whether natural selection has been operating and still operates in modern humans—and at what rate—has been the subject of much debate. Until recently, it was often held that human evolution had come to an end about 40,000 to 50,000 years ago (see, e.g., ref. (1)). However, new evidence that has been accumulating over the last decade suggests that natural selection has been operating in humans over the past few thousand years (2–4) and that a number of adaptations—such as lactase persistence (5), resistance to malaria (6), and adaptation to high altitude (7)—have occurred relatively recently. It has also been shown that height and body mass index have been under selection in Europeans (8).

In parallel, a number of recent studies have sought to examine the association between lifetime reproductive success (LRS)—the number of children an individual ever gave birth to or fathered—and various phenotypes in contemporary human populations. (In modern populations with low mortality, fitness can be reasonably approximated by LRS (9, 10), notwithstanding some caveats discussed below.) These studies have typically found that natural selection has been operating in contemporary humans (9, 11–14). It has also been shown that there was significant variance in relative fitness in a preindustrial human population, such that there was much potential for natural selection (15).

However, this literature has analyzed the relationship between *phenotypes* and LRS, and natural selection occurs only when genotypes that are associated with the phenotypes covary with reproductive success. This literature’s conclusions regarding ongoing natural selection are thus particularly sensitive to assumptions that are needed to estimate the relationship between genotypes and phenotypes and to the inclusion in the analysis of all correlated phenotypes with causal effects on fitness (16, 17).^*^ Some of those assumptions have been criticized and debated (e.g., (18)), and it has proven challenging to include all relevant correlated phenotypes in analyses of selection in natural populations (17).

Recent advances in molecular genetics now make it possible to look directly at the relationship between LRS and genetic variants associated with various phenotypes, thus eliminating those potential confounds. Here, I examine the association between relative LRS (rLRS)—the ratio of LRS to the mean LRS of individuals of the same gender born in the same years—and genetic variants associated with various phenotypes, for a sample of females and males in the Health and Retirement Study (HRS). Using rLRS instead of LRS as the measure of fitness helps control for the effects of time trends in LRS and makes it possible to interpret my estimates as rates of natural selection (9, 16), as I discuss below. (My results are robust to using LRS instead of rLRS as the measure of fitness.)

The phenotypes I analyze are body mass index (BMI), educational attainment (EA), fasting glucose concentration (GLU), height (HGT), schizophrenia (SCZ), plasma concentrations of total cholesterol (TC), and age at menarche (AAM; in females). These phenotypes were selected on the basis of previous evidence showing that selection acts on some of them (see, e.g., (9)) and because summary statistics (i.e., the estimated effects of the single nucleotide polymorphisms (SNPs) on the phenotypes) from previous large-scale genome-wide association studies (GWAS) are available for them (19–25).

The HRS is a representative longitudinal panel study of ~20,000 Americans shortly before and during retirement. It is well suited for this study for several reasons. First, the HRS was designed to be representative of the US population over the age of 50 (26), which makes it possible to generalize my results to the entire US population of European ancestry born in the years of my study sample. In addition, individuals in the study are in the later stages of their lives, when they have typically completed their lifetime reproduction. Nonetheless, as I discuss below, selection bias due to incomplete genotyping of the study participants and differential survival remains a concern, although genotyped individuals do not appear to differ markedly from non-genotyped individuals in my study sample.

To mitigate the risks of confounding by population stratification, my analyses focus on unrelated individuals of European ancestry and control for the top 20 principal components of the genetic relatedness matrix (which capture the main dimensions along which the ancestry of the individuals in the dataset vary (27)). To mitigate the risk of selection bias due to differential mortality, and to ensure that the LRS variable is a good proxy for completed fertility, I limit my analyses to individuals born between 1931 and 1953 and who were at least 45 years old (for females) or 50 years old (for males) when asked the number of children they ever gave birth to or fathered. I refer to the resulting sample as the “study sample.” I performed my main analyses separately for females and males, as different selection gradients can operate across genders.

In a recent paper, Tropf et al. (28) used genetic data to study the relationship between LRS and age at first birth in a sample of females, and found that the two phenotypes are negatively genetically correlated. My analyses complement theirs in several important ways: my analyses cover both females and males and seven different phenotypes; they include childless individuals (who can have an important impact on the gene pool by foregoing reproduction); they indirectly leverage the statistical power of previous large-scale GWAS to estimate the relationship between rLRS and genetic variants associated with the phenotypes, thus increasing the precision of my estimates for some phenotypes;^†^ and I translate selected estimates into interpretable measures of the rate at which natural selection has been operating.

## Phenotypic Evidence for Natural Selection

I begin by looking at the phenotypic evidence for natural selection in the HRS. The HRS contains phenotypic variables for four of the seven phenotypes I study: BMI, EA, HGT, and TC. (The phenotypic variable for TC is an indicator for a self-reported health problem with high cholesterol, and not plasma concentrations of total cholesterol as in the GWAS of TC.) Table S1 reports summary statistics for these and for the other phenotypic variables I use, both for all individuals in the study sample and for the genotyped individuals in the study sample. As can be seen, the two samples look remarkably similar. Table 1 reports estimates from separate regressions of rLRS on each of these phenotypic variables (and on control variables) for the sample of all individuals (genotyped and not genotyped) in the study sample. Stouter females and males, less educated females and males, and smaller females have significantly higher rLRS (*P* ≤ 0.001 in all cases). The estimates for the sample of genotyped individuals are very similar (Table S2), thus suggesting that the two samples are similar in terms of the selection gradients that were operating on the various phenotypes.

**Table 1.**
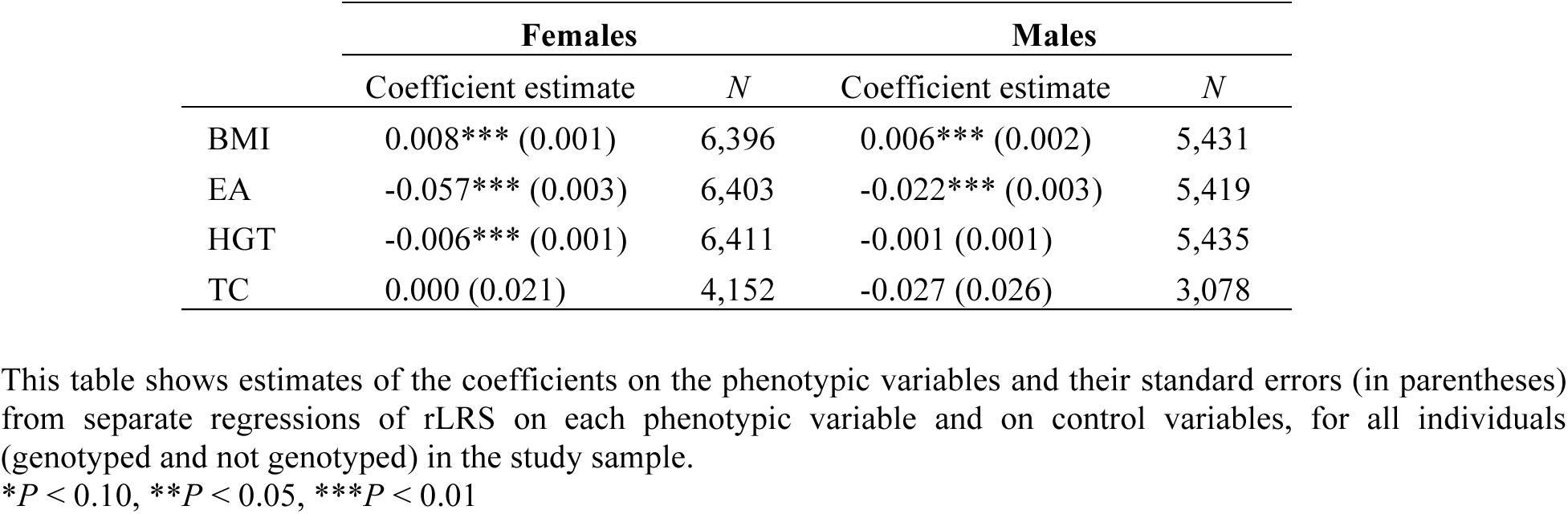
Estimates from separate regressions of rLRS on each phenotypic variable, for all individuals

|  | Females |  | Males |  |
| --- | --- | --- | --- | --- |
|  | Coefficient estimate | *N* | Coefficient estimate | *N* |
| BMI | 0.008*** (0.001) | 6,396 | 0.006*** (0.002) | 5,431 |
| EA | -0.057*** (0.003) | 6,403 | -0.022*** (0.003) | 5,419 |
| HGT | -0.006*** (0.001) | 6,411 | -0.001 (0.001) | 5,435 |
| TC | 0.000 (0.021) | 4,152 | -0.027 (0.026) | 3,078 |

This table shows estimates of the coefficients on the phenotypic variables and their standard errors (in parentheses) from separate regressions of rLRS on each phenotypic variable and on control variables, for all individuals (genotyped and not genotyped) in the study sample.
\* $P < 0.10$ , \*\* $P < 0.05$ , \*\*\* $P < 0.01$

As mentioned, without assumptions to estimate the relationship between genotypes and phenotypes and without considering all correlated phenotypes with possible causal effects on fitness, it is not possible to translate these estimates into estimates of evolutionary change—even over a single generation. Previous research, however, has established that these phenotypes are all moderately to highly heritable (29); notwithstanding the possible effects of correlated phenotypes, this is suggestive that genotypes associated with BMI, EA, and HGT covary with fitness and that natural selection has been operating on these phenotypes.

## Genetic Evidence for Natural Selection

To test directly whether natural selection has been operating on the genetic variants associated with BMI, EA, GLU, HGT, SCZ, TC, and AAM, the summary statistics from the latest GWAS of these phenotypes were used to construct polygenic scores that partially predict the genotyped individuals’ phenotypes based on their genotyped SNPs. To avoid overfitting (30), the GWAS summary statistics used are all based on meta-analyses that exclude the HRS. For the main analyses, LDpred (31) was employed to construct the scores. LDpred uses a prior on the SNPs’ effect sizes and adjusts summary statistics for linkage disequilibrium (LD) between SNPs to produce scores that have higher predictive power than the alternatives. (My results are robust to using scores constructed with PLINK (32), which does not adjust the summary statistics for LD between SNPs). The scores were standardized to have mean zero and a standard deviation of one. Additional details on the construction of the scores are provided in *Materials and Methods* and in *Supporting Information*.

Fig. 1 shows the highest previously reported *R*^*2*^ of the scores from the articles reporting the GWAS whose summary statistics were used to construct the scores, as well as the incremental *R*^*2*^ of the scores of BMI, EA, HGT, and TC (for which there are phenotypic variables in the HRS) in the study sample. (The incremental *R*^*2*^ of the score of a phenotype is defined as the difference between the *R*^*2*^ of the regression of the phenotype on controls for gender and birth year, the top 20 principal components of the genetic relatedness matrix, and the score, and the *R*^*2*^ of the same regression but without the score.) The incremental *R*^*2*^ range from 0.012 (for TC) to 0.174 (for HGT) and are all significantly larger than zero; nonetheless, they are all much smaller than estimates of the phenotypes’ heritability in the existing literature (29), implying that the scores are very imperfect proxies for the individuals’ true genetic scores (defined as the sum of the true average causal effects of all their alleles) for the various phenotypes.^℡^

**Fig. 1.**
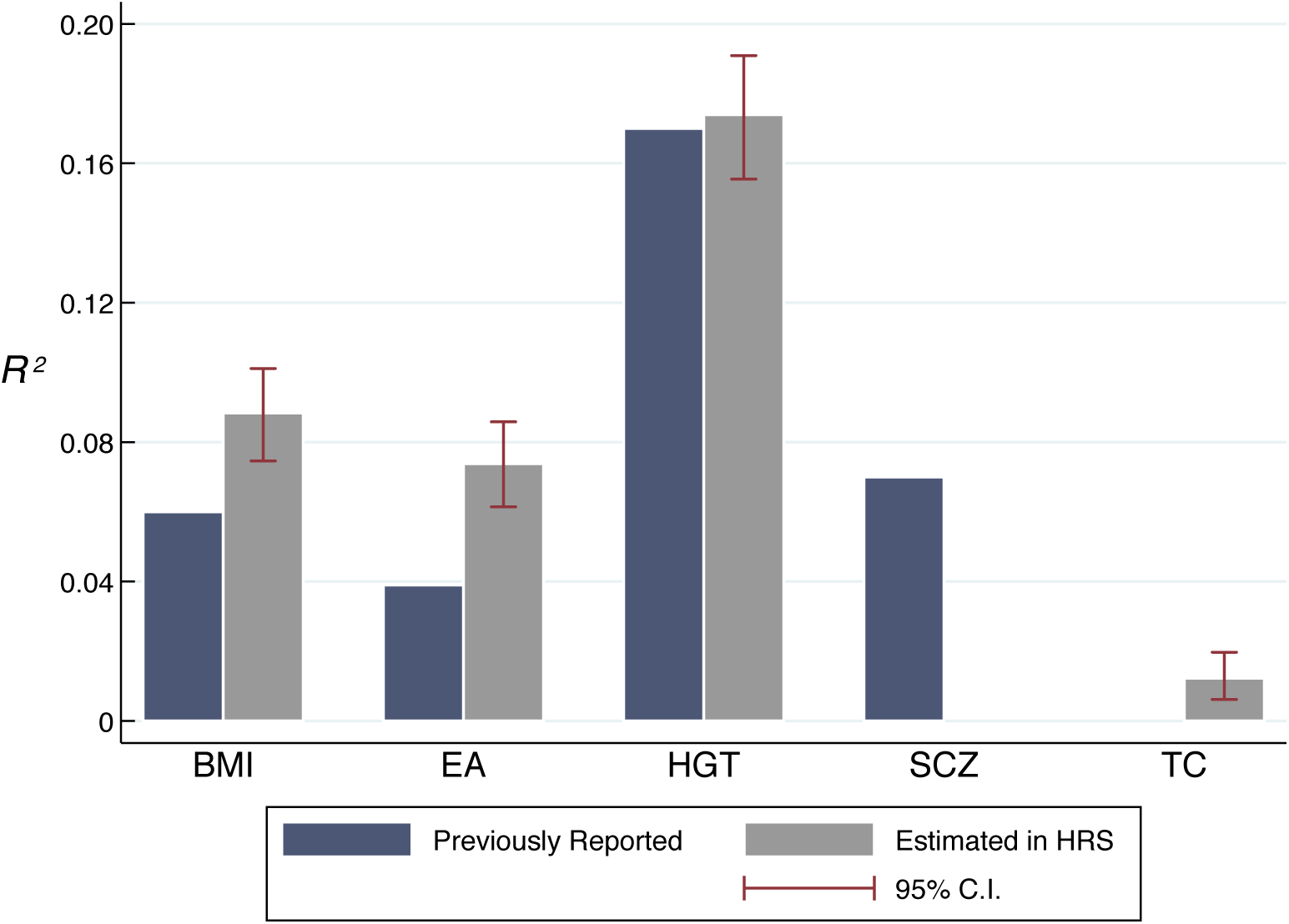
Predictive power (*R*^2^) of polygenic scores of the various phenotypes. “Previously reported”: highest previously reported *R*^*2*^ of scores from prediction analyses from the articles reporting the GWAS whose summary statistics were used to construct the scores for each phenotype. The previously reported *R*^*2*^ for SCZ is the *R*^*2*^ on the liability scale; the *R*^*2*^ of the score was not reported for GLU, AAM, and TC. “Estimated in HRS”: estimates of the incremental *R*^*2*^ of the LDpred scores used in this article, with percentile confidence intervals estimated with the nonparametric bootstrap with 1,000 bootstrap samples.

Table 2 reports estimates from separate regressions of rLRS on the polygenic scores of the various phenotypes (and on control variables, which include the top 20 principal components of the genetic relatedness matrix (27)). The score of EA is significantly negatively associated with rLRS for both females (*P* = 0.002) and males (*P* = 0.013). The association remains significant after Bonferroni correction for 13 tests (the number of estimates reported in Table 2) for females, but not for males (Bonferroni-corrected *P* = 0.022 for females, = 0.174 for males). The estimates for females and males are similar, and the association is also significant in the sample of females and males together (Table S3, *P* = 1.2 × 10^-5^) and remains significant after Bonferroni correction for 7 tests (the number of phenotypes) (Bonferroni-corrected *P* = 8.3 × 10^-5^). Fig. 2 shows the mean polygenic score of EA as a function of LRS, by sex. Both females and males who had no children have a significantly higher mean score of EA than those who had one or more children (*P* < 0.005 in both cases, unpaired *t*-tests). Thus, the negative association between rLRS and the score of EA appears to be driven primarily by score differences between individuals with and without children.

**Fig. 2.**
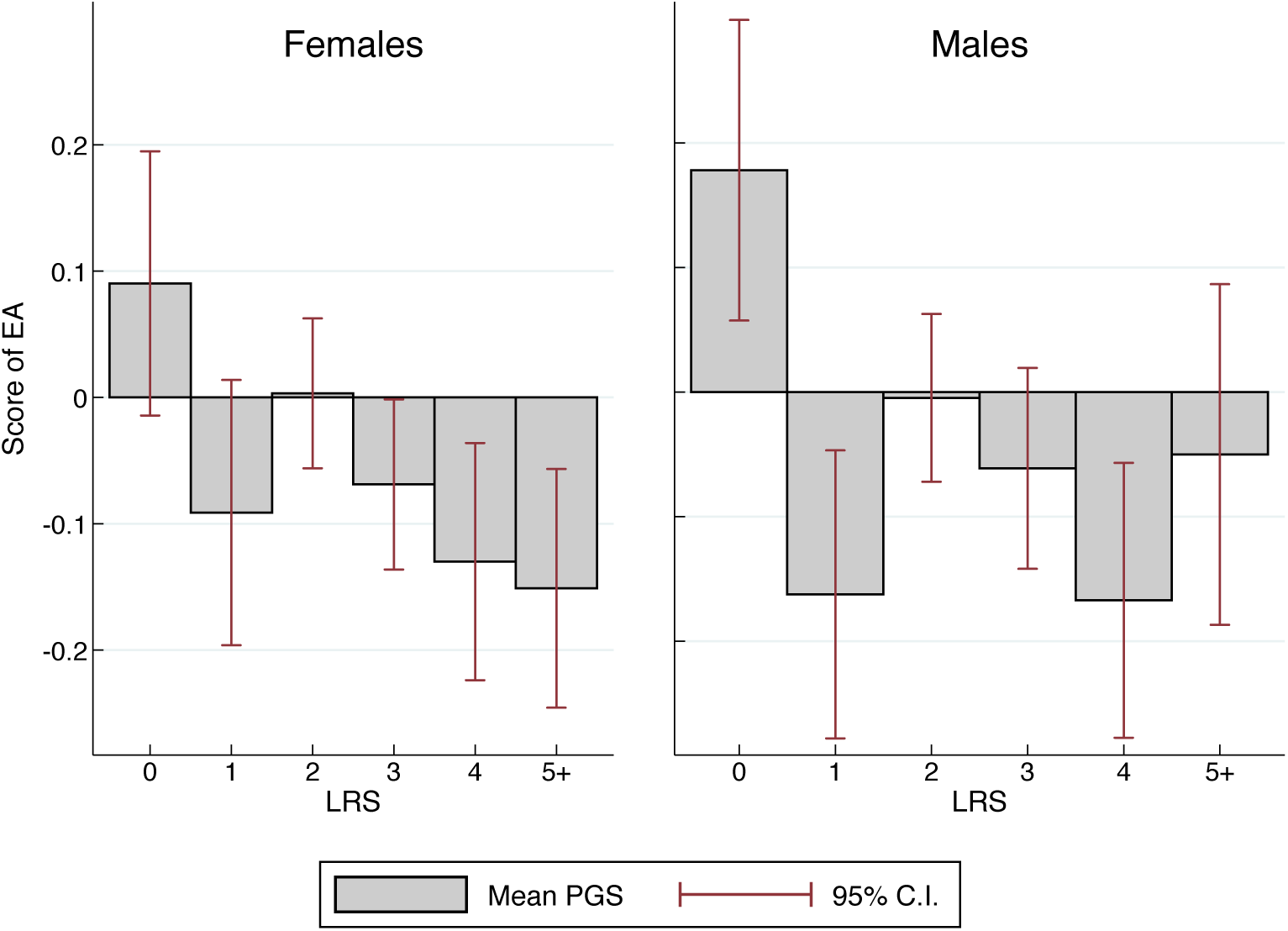
Mean polygenic score of EA as a function of LRS, for females and males in the study sample.

**Table 2.**
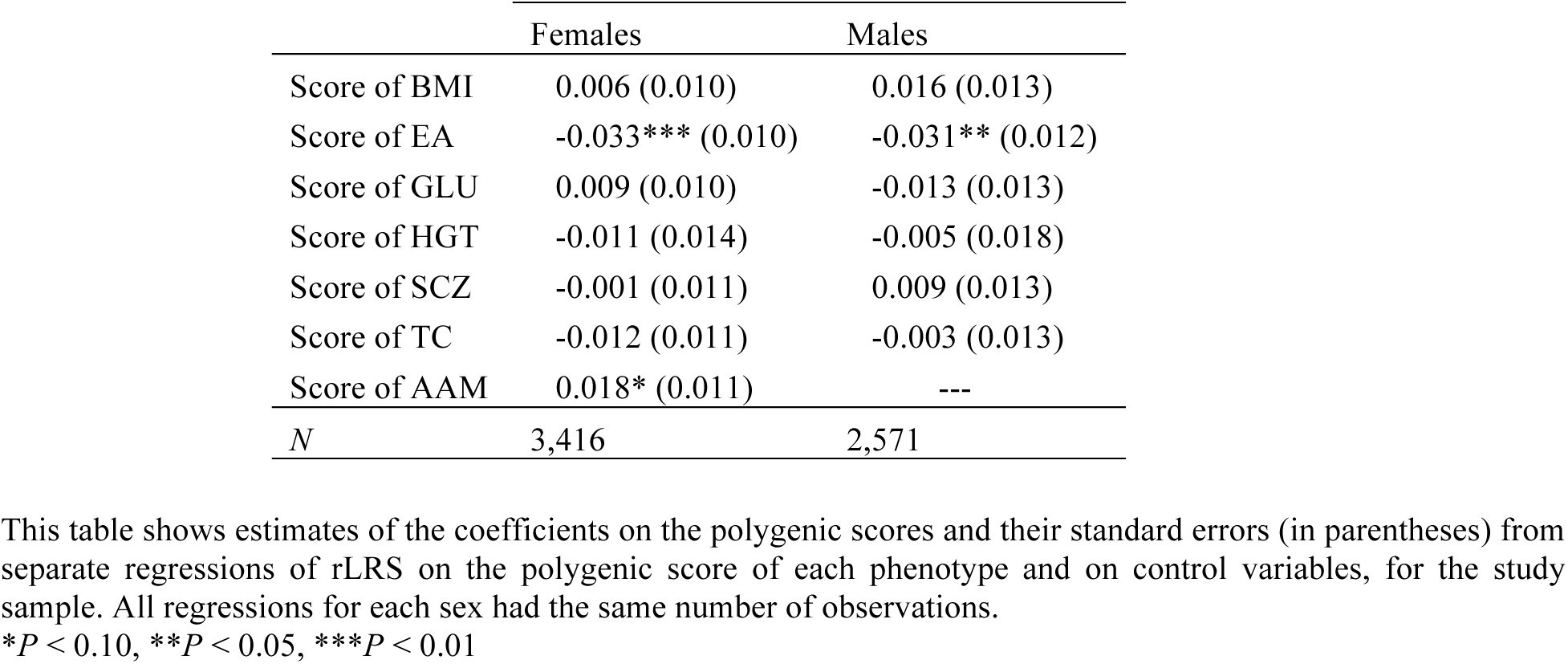
Estimates from separate regressions of rLRS on the polygenic score of each phenotype

|  | Females | Males |
| --- | --- | --- |
| Score of BMI | 0.006 (0.010) | 0.016 (0.013) |
| Score of EA | -0.033*** (0.010) | -0.031** (0.012) |
| Score of GLU | 0.009 (0.010) | -0.013 (0.013) |
| Score of HGT | -0.011 (0.014) | -0.005 (0.018) |
| Score of SCZ | -0.001 (0.011) | 0.009 (0.013) |
| Score of TC | -0.012 (0.011) | -0.003 (0.013) |
| Score of AAM | 0.018* (0.011) | --- |
| *N* | 3,416 | 2,571 |

This table shows estimates of the coefficients on the polygenic scores and their standard errors (in parentheses) from separate regressions of rLRS on the polygenic score of each phenotype and on control variables, for the study sample. All regressions for each sex had the same number of observations.
\* $P < 0.10$ , \*\* $P < 0.05$ , \*\*\* $P < 0.01$

The polygenic score of AAM is also significantly and positively associated with rLRS for females at the 10% level (*P* = 0.080), but this association does not remain significant after Bonferroni correction for 13 tests. I therefore interpret it as being weakly suggestive that genetic variants associated with higher AAM may have been selected for. The polygenic scores of the other phenotypes (BMI, GLU, HGT, SCZ, TC) are not robustly significantly associated with rLRS. Although these estimates are small in magnitude and insignificant, this could be because my polygenic scores are imperfect proxies for the true genetic scores and does not prove that natural selection has not been operating on genetic variants associated with those phenotypes.

According to the Robertson-Price identity (33, 34), the directional selection differential of a “character” is equal to the genetic covariance between the character and relative fitness. (A character is an observable feature of an organism, and its directional selection differential is the change in its mean value due to natural selection in one generation.) As I show in *Supporting Information*, if we define the polygenic scores as the characters of interest, it follows that the coefficients on the scores reported in Table 2 can be interpreted as the directional selection differentials of the scores themselves, expressed in Haldanes—i.e., each coefficient equals the implied change in the score that will occur due to natural selection in one generation, expressed in standard deviations of the score per generation. Hence, the estimates from Table 2 imply that natural selection has been operating on the score of EA at a rate of -0.033 Haldanes among females and of -0.031 Haldanes among males in the study sample. (Even if the mechanism that underlies the negative association between rLRS and the score of EA is that more educated people *choose* to have fewer children, it would still be the case that natural selection has been operating.)

I rescaled these estimates of the directional selection differential of the score of EA to express them in years of education per generation rather than in Haldanes (*Supporting Information*). My rescaled estimates imply that natural selection has been operating on the score of EA at rates of - 0.022 (95% CI: -0.036 to -0.009) and -0.022 (95% CI: -0.040 to -0.004) years of education per generation for females and males—or about minus one week of education per generation for both sexes.

I also obtained estimates of the directional selection differential of EA (or, equivalently, of the true genetic score of EA), which is equal (under some assumptions) to the directional selection differential of the polygenic score of EA multiplied by the ratio of the heritability of EA to the *R*^*2*^ of the score of EA (*Supporting Information*). (I assume the heritability of EA to be 0.40, based on a recent meta-analysis of existing heritability estimates of EA (35).) Generalizing my results from the study sample to the general population, my estimates imply that natural selection has been operating on EA at rates of -1.30 (95% CI: -2.12 to -0.54) and -1.53 (95% CI: -2.85 to -0.31) months of education per generation among US females and males of European ancestry born between 1931 and 1953. As I discuss below, these rates are small relative to the secular increases in EA that have been observed over the past few generations.

I performed a number of checks to verify the robustness of my results. First, I repeated the analyses with LRS instead of rLRS. Second, I used polygenic scores constructed with PLINK (32) instead of LDpred. Third, I only included individuals aged no more than 70 in 2008 (the last year in which individuals were genotyped) and at least 50 years old (for females) or 55 years old (for males) when asked their number of children—to mitigate the risk of selection bias due to differential mortality and to ensure that almost every individual had completed fertility when asked his or her number of children. Fourth, I included the HRS0 cohort of individuals born between 1924 and 1930 together with the study sample. (As detailed in *Materials and Methods*, I define cohorts based on the individuals’ birth years; the study sample includes the HRS1, HRS2, and HRS3 cohorts, but excludes the HRS0 cohort because of possible selection bias based on mortality.) Table S4 presents the results of those checks. In all cases, the results for EA are robust. Further, for the score of EA for both females and males and for the score of AAM (for females), the estimates are not significantly different from one another at the 5% level across the HRS1, HRS2, and HRS3 cohorts (Table S3 and *t*-tests of the interactions between the coefficients on the scores and cohort dummies).

Following Lande and Arnold (16), I also estimated quadratic regressions of rLRS on all the polygenic scores and their squares and interactions together and on control variables, to test for nonlinear selection (Table S5 and *Supporting Information*). I found no convincing evidence that nonlinear selection has been operating on the genetic variants associated with the various phenotypes.

## Discussion

My results suggest that natural selection has been operating on the genetic variants associated with EA, and possibly with AAM. Though I find no evidence that natural selection has been operating on the genetic variants associated with the other phenotypes or that nonlinear selection has been operating, I emphasize that this could be because my polygenic scores are imperfect proxies for the true genetic scores, which limits the statistical power of my analyses.

My estimates of the negative associations between rLRS and both phenotypic EA and the polygenic score of EA are consistent with previous findings of negative associations between LRS and phenotypic EA in samples of females (36–39), males (36, 39), and females and males together (40) in contemporary Western populations, though positive phenotypic associations have also been reported for males (37). Conley et al. (41) report a negative phenotypic association for females and males together, as well as a negative correlation between LRS and a score of EA (constructed with the summary statistics from an earlier, smaller GWAS of EA). To my knowledge, few articles have investigated the relationship between phenotypic AAM and LRS in contemporary Western populations. Kirk et al. (42) find a quadratic relationship that is suggestive of stabilizing selection, but find no genetic covariation (using behavioral genetic techniques in a sample of twins). Consistent with the results of my regressions of rLRS on phenotypic BMI and HGT (Table 1), there is previous phenotypic evidence of positive selection for weight and negative selection for HGT in females (9, 43). Previous studies have also documented a positive (9) and an inverted-U (43) relationship between LRS and phenotypic HGT in males, and have found evidence of negative selection for TC and of stabilizing selection for GLU in females (11), in contemporary Western human populations.

Consistent with the results from previous studies with phenotypic data (e.g., (9)), my results suggest that natural selection has been operating slowly relative to the rapid secular changes that have occurred over the past few generations, presumably due to cultural and environmental factors. For instance, my estimate of a directional selection differential of EA of about -1.5 months of education per generation pales in comparison with the increase of 6.2 years in the mean level of EA that took place for native-born Americans born between 1876 and 1951 (44) (which is equivalent to about 2 years of education per generation). Moreover, although I find suggestive evidence that genetic variants associated with higher AAM may have been selected for, AAM has substantially *decreased* in contemporary Western populations (45). And although I find no evidence of selection for the genetic variants associated with BMI and HGT, both phenotypes have markedly increased over the past century (46). Thus, although natural selection is still operating, the environment appears to have achieved an “evolutionary override” (28) on the measurable phenotypes I study.

As shown in Okbay et al. (20), the association between the score of EA and EA is not likely to be driven by the effects of culture, the environment, or population stratification, and is likely to reflect the true causal effects of multiple genetic variants. For instance, in cohorts that are independent of those used in the GWAS of EA, the score remains significant in regressions of EA on the score when family fixed effects are also included. Moreover, estimates from an LD score regression (47)—which disentangles the signal due to the genetic variants’ causal effects from the signal due to confounding biases—suggest that stratification is not a major source of bias in the GWAS summary statistics of EA. Okbay et al. also analyzed the summary statistics of EA and obtained sizeable and significant estimates of the genetic correlation between EA and several neuropsychiatric and cognitive phenotypes, as well as of the genetic variance of EA accounted for by SNPs annotated to the central nervous system relative to other SNPs. Thus, while it is not possible to rule out with certainty that my results are (at least partly) confounded by stratification, stratification is unlikely to be an important concern.

Several additional caveats should be kept in mind when interpreting my results. First, rLRS is not a perfect proxy for long-term genetic contribution. Among other possible reasons for this, a tradeoff between the quantity and quality of children has been documented in preindustrial human societies and may still exist in modern societies (48). In the presence of such a tradeoff, the number of grandchildren or third-generation descendants is a better measure of fitness— though most datasets (including the HRS) lack such data, and it has been shown that LRS and number of grand-offspring were perfectly genetically correlated in a post-industrial human population (10). Also, in growing populations, individuals who successfully reproduce earlier in life tend to have higher fitness (49), but rLRS does not account for fertility timing. In the case of EA, individuals with high EA typically have children at a more advanced age, which may further reduce their fitness. Alternative measures of fitness—such as the intrinsic rate of increase (the exponentiated Lotka’s *r*)—account for fertility timing, but require data on the age at birth of every offspring and do not always perform better in natural populations (50). A second caveat is that it is not possible to translate my estimates into projected evolutionary changes over more than one generation, because my results do not account for the effects of all phenotypes that correlate genetically with the phenotypes I study and that also have causal effects on fitness (16).^§^ Furthermore, since the cultural environment changes through time, the selection gradients that existed from 1931 to 1953 may not apply to earlier and subsequent periods, which makes long-term projections problematic. For instance, it has been shown that the demographic transition has significantly changed the selective forces in some populations (39, 51–53).

Lastly, there are several reasons why my results in the study sample of genotyped individuals might not be fully generalizable to the entire US population of European ancestry born between 1931 and 1953. First, the HRS only targets individuals who survived until age 50, and about 10% of female and 15% of male Americans born in 1940 died before reaching age 50, based on data from the United States Social Security Administration (54). Second, in the study sample, only 85% of the participants were still alive in 2008 (the last year when they could be genotyped), 69% were asked to be genotyped, and 59% consented to be genotyped. That being said, a comparison of the summary statistics for all individuals in the study sample and for the genotyped individuals in the study sample (Table S1) suggests that there are no important differences between the two samples, and the results of the phenotypic regressions are very similar across the two samples (Tables 1 and S4).

In sum, while keeping those limitations in mind, my results strongly suggest that genetic variants associated with EA have slowly been selected against among both female and male Americans of European ancestry born between 1931 and 1953, and that natural selection has thus been occurring in that population—albeit at a rate that pales in comparison with the rapid secular changes that have occurred in recent generations. My results also suggest that genetic variants positively associated with AAM may have been positively selected for among females in that population. As larger GWAS are conducted and better estimates of genetic variants’ effects on various phenotypes become available, polygenic scores will become more precise. The eventual completion of a GWAS of LRS will also make it possible to use other methods, such as LD score regressions (55), to estimate the genetic covariance between LRS and other phenotypes. Future studies that address the above-mentioned limitations will be able to leverage these developments to replicate my results and to obtain more precise estimates of the rate at which natural selection has been and is occurring in humans.

## Materials and Methods

### The Study Sample and the Cohorts

The Health and Retirement Study (HRS) is a longitudinal panel study for which a representative sample of approximately 20,000 Americans have been surveyed every two years since 1992. My main analyses focus on individuals born between 1931 and 1953. To reduce the risks of confounding by population stratification, I restrict the analyses to unrelated individuals of European ancestry (i.e., non-Hispanic White individuals). To ensure that the lifetime reproductive success (LRS) variable is a good proxy for completed fertility, I only include females who were at least 45 years old and males who were at least 50 years old when asked the number of children they ever gave birth to or fathered. Further, to ensure that the sample of individuals who have been successfully genotyped (whose DNA samples were collected between 2006 and 2008) is comparable to the sample of individuals who have not, I only include individuals who were enrolled in the HRS and asked the number of children they ever gave birth to or fathered in 2008 or earlier. This left 6,414 females and 5,436 males with phenotypic data and 3,416 unrelated females and 2,571 unrelated males who have been successfully genotyped and who passed the quality control filters described in *Supporting Information* (and for whom I could thus construct polygenic scores). I refer to the resulting sample as the “study sample.”

For some specifications, I divided the study sample into three non-overlapping cohorts based on the individuals’ birth years. This allowed me to test the robustness of my results across cohorts (my definition of the cohorts resembles the HRS’ definition, which recruited its different cohorts at different times). Table S6 summarizes the three cohorts—which I label HRS1 (birth years 1931 to 1941), HRS2 (birth years 1942 to 1947), and HRS3 (birth years 1948 to 1953)—as well as the HRS0 cohort of individuals born between 1924 and 1930. To mitigate the risk of selection bias based on mortality, I excluded the HRS0 cohort of individuals born between 1924 and 1930 from the study sample (Table S6 and *Supporting Information*), but my main results are robust to the inclusion of that cohort (Table S4). For the same reason, I excluded individuals born prior to 1924 from the study sample. I also excluded individuals born after 1953 from the study sample, as very few of them have been genotyped.

### Phenotypic Variables

For my baseline analyses, I operationalize relative fitness with the relative lifetime reproductive success (rLRS) variable. As Fig. S1 shows, LRS for females and males declined gradually between 1931 and 1953, from around 3 children in the early 1930s to 2 children around 1950. Table S1 presents summary statistics for birth year, LRS, and childlessness, as well as for the phenotypic variables for body mass index (BMI), educational attainment (EA), height (HGT), and total cholesterol (TC). *Supporting Information* provides details on the construction of these variables. The HRS does not contain phenotypic variables for fasting glucose concentration (GLU), schizophrenia (SCZ), and age at menarche (AAM; in females).

### Quality Control of the Genotypic Data and Polygenic Scores

I followed the HRS recommendations regarding the use of the genotypic data (“Quality Control Report for Genotypic Data”). The individuals’ genotyped single nucleotide polymorphisms (SNPs) that passed the quality control filters and that were present in the phenotypes’ GWAS summary statistics files were used to construct the polygenic scores. Depending on the phenotype, there were between 505,254 and 544,493 such overlapping SNPs (except for GLU, for which there were only 22,895 such overlapping SNPs). The average sample sizes across the SNPs used to construct the scores are 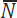_*BMI*_ = 232,186, 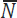_*EA*_ = 386,098, 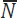_*HGT*_ = 243,630, and 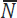_*TC*_ = 92,793 individuals; the summary statistics for GLU, SCZ, and MEN did not contain sample size information, but the reported samples sizes for the main GWAS of these phenotypes are *N*_*GLU*_ = 133,010, *N*_*scz*_ ≈ 80,000, and *N*_*men*_ = 132,989 individuals. The GWAS summary statistics used to construct the scores are all based on meta-analyses that exclude the HRS.

For the main analysis, I used LDpred (31) to construct the polygenic scores; for a robustness check, I also constructed polygenic scores with PLINK (32). (The polygenic scores of EA were constructed and provided to me by the Social Science Genetic Association Consortium (SSGAC), following the procedure described here and which I used to construct the other scores.) Both the LDpred and the PLINK scores for an individual are weighted sums of the individual’s genotype across all SNPs. For the PLINK scores, the weight for each SNP is the GWAS estimate of the SNP’s effect, which captures the causal effects of both the SNP and of SNPs that are in linkage disequilibrium (LD); for the LDpred scores, the weight for each SNP is the LDpred estimate of the SNP’s causal effect, which LDpred calculates by adjusting the SNPs’ GWAS estimates with a prior on the SNPs’ effect sizes and information on the LD between the SNPs from a reference panel. The LDpred prior on the SNPs’ effect sizes depends on an assumed Gaussian mixture weight. For each phenotype, LDpred scores were constructed for a range of Gaussian mixture weights, and I selected the score with the weight that maximizes the incremental *R*^*2*^ of the score or the correlations between the score and known correlates of the phenotype. Both the LDpred and PLINK scores were standardized to have mean zero and a standard deviation of one. *Supporting Information* provides more information on the quality control steps and the construction of the polygenic scores, and Table S7 shows the parameters used to construct the scores and the sources for each phenotype’s summary statistics.

### Association Analyses

For each of BMI, EA, HGT, and TC—for phenotypic variables are available in the HRS—I regressed rLRS on the corresponding phenotypic variable, separately for females and males; those regressions included birth year dummies and HRS-defined cohort dummies and were estimated by ordinary least squares (OLS). For all phenotypes, I also regressed rLRS on the polygenic score of the phenotype in various samples; the regressions also included birth year dummies, HRS-defined cohort dummies, and the top 20 principal components of the genetic relatedness matrix (to control for population stratification (27); see also Section 5 of the Supplemental Material of ref. (56)), and were also estimated by OLS. For the regressions in the sample of females and males together, I also controlled for sex and only included the respondent with the lowest person number (PN, an HRS identifier) in each household, as spouses very often have the same number of children, which induces a complex correlation structure between the error terms (the results for the score of EA are robust to alternative ways of selecting one respondent per household). In all results tables, I report the coefficient estimates and standard errors, with stars to indicate statistical significance; *P*-values are included in the log files available on my website. The standard errors and the P-values implied by the stars in the tables reporting my estimates from regressions of rLRS on the LDpred scores do not account for the uncertainty stemming from the selection of the Gaussian mixture weights for the LDpred scores; however, the fact that my results are robust to the use of the PLINK scores instead of the selected LDpred scores implies that my main results are not driven by this weight selection procedure.

### Directional Selection Differentials

Based on the Robertson-Price identity (33, 34), the directional selection differential of a character is equal to its genetic covariance with relative fitness. As I show in *Supporting Information*, it follows that the estimates of the coefficients on the polygenic scores reported in Table 2 can be interpreted as directional selection differentials of the scores, expressed in Haldanes (one Haldane is one standard deviation per generation). *Supporting Information* also shows how to rescale the estimates of the directional selection differential for the score of EA to express them in years of education per generation instead of in Haldanes, and shows how to obtain estimates of the directional selection differential of EA (or, equivalently, of the true genetic score of EA—rather than of the polygenic score of EA) expressed in years of education per generation (under some assumptions). I used the nonparametric bootstrap method with 1,000 bootstrap samples to obtain percentile confidence intervals for the estimates of the directional selection differentials.

## Footnotes

* For instance, if phenotypes 1 and 2 are phenotypically but not genetically correlated and if only phenotype 2 is under selection, an analysis based on phenotypic data that does not include phenotype 2 or that fails to account for the lack of genetic correlation may erroneously conclude that phenotype 1 is also under selection.

‡ An alternative to my “score regression” approach (described below) is the bivariate GREML method (57) (used by Tropf et al.), but it is not well-suited for the present study. It has very low power and yields imprecise estimates of the genetic correlation in samples of moderate size like the HRS, given the low SNP heritability of rLRS (28) (according to the GCTA-GREML Power Calculator (58)). It also requires a dataset with phenotypic data for every studied phenotype and assumes normally distributed phenotypes (which is not realistic for rLRS).

‡ The *R*^*2*^ of the polygenic score of a phenotype is bounded by the phenotype’s heritability (30) and depends in part on the precision with which the effects of the individual genetic variants were estimated in the GWAS of that phenotype, which in turn depends on the GWAS sample size. Future, larger GWAS should allow more precise estimation of the effects of the genetic variants and the construction of more precise scores.

§ Selection on phenotypes that are genetically correlated with the phenotypes of interest impacts their genetic covariance, which in turn impacts the selection gradients on the phenotypes of interest in future generations.

## Acknowledgments

I thank David Cesarini, Joseph Henrich, Lawrence Katz, Iain Mathieson, Steven Pinker, Alkes Price, Stephen Stearns, and Peter Visscher for helpful comments. I also thank Dan Benjamin and David Laibson for helpful comments and postdoctoral supervision. The polygenic scores of EA were constructed and provided by the Social Science Genetic Association Consortium (SSGAC); I thank Aysu Okbay for constructing these scores on behalf of the SSGAC.

